# Generation of anisotropic strain dysregulates wild-type cell division at the interface between host and oncogenic tissue

**DOI:** 10.1101/578146

**Authors:** Megan Moruzzi, Alexander Nestor-Bergmann, Georgina K. Goddard, Keith Brennan, Sarah Woolner

## Abstract

Epithelial tissues are highly sensitive to anisotropies in mechanical force, with cells altering fundamental behaviours such as cell adhesion, migration and cell division [1-5]. It is well known that in the later stages of carcinoma (epithelial cancer), the presence of tumours alters the mechanical properties of a host tissue and that these changes contribute to disease progression [6-9]. However, in the earliest stages of carcinoma, when a clonal cluster of oncogene-expressing cells first establishes in the epithelium, the extent to which mechanical changes alter cell behaviour in the tissue as a whole remains unclear. This is despite knowledge that many common oncogenes, such as oncogenic Ras, alter cell stiffness and contractility [10-13]. Here, we investigate how mechanical changes at the cellular level of an oncogenic cluster can translate into the generation of anisotropic strain across an epithelium, altering cell behaviour in neighbouring host tissue. We generated clusters of oncogene-expressing cells within otherwise normal *in vivo* epithelium, using *Xenopus laevis* embryos. We find that cells in *k*Ras^V12^, but not *c*MYC, clusters have increased contractility, which introduces radial stress in the tissue and deforms surrounding host cells. The strain imposed by *k*Ras^V12^ clusters leads to increased cell division and altered division orientation in the neighbouring host tissue, effects that can be rescued by reducing actomyosin contractility specifically in the *k*Ras^V12^ cells. Our findings indicate that some oncogenes can alter the mechanical and proliferative properties of host tissue from the very earliest stages of cancer development, changes which have the potential to contribute to tumorigenesis.

## RESULTS AND DISCUSSION

### Modelling early stage carcinoma in *Xenopus laevis*

To investigate how mechanical changes might alter cell behaviour in a model of early stage carcinoma, we chose two common oncogenes: *k*Ras^V12^ and *c*MYC, which we hypothesized would have differing effects on tissue mechanics. Ras GTPases are well-known to sit upstream of actomyosin contractility and previous studies in single cells have shown that constitutively active Ras mutations alter cell stiffness [11, 12]. In contrast, there is little evidence that *c*MYC would have a direct effect on the mechanical properties of cells or tissues. To produce a cluster of oncogene-expressing cells within wild-type *in vivo* epithelial tissue, GFP-*k*Ras^V12^ or GFP-*c*MYC mRNA was injected into a single cell of a stage 6 (32 cell) *Xenopus laevis* embryo (Figure 1A). By early gastrula stage, a cluster of GFP-expressing cells consistently developed in the superficial layer of the animal cap epithelium (Figures 1B-D, Figure S1A-B).

**Figure 1:**
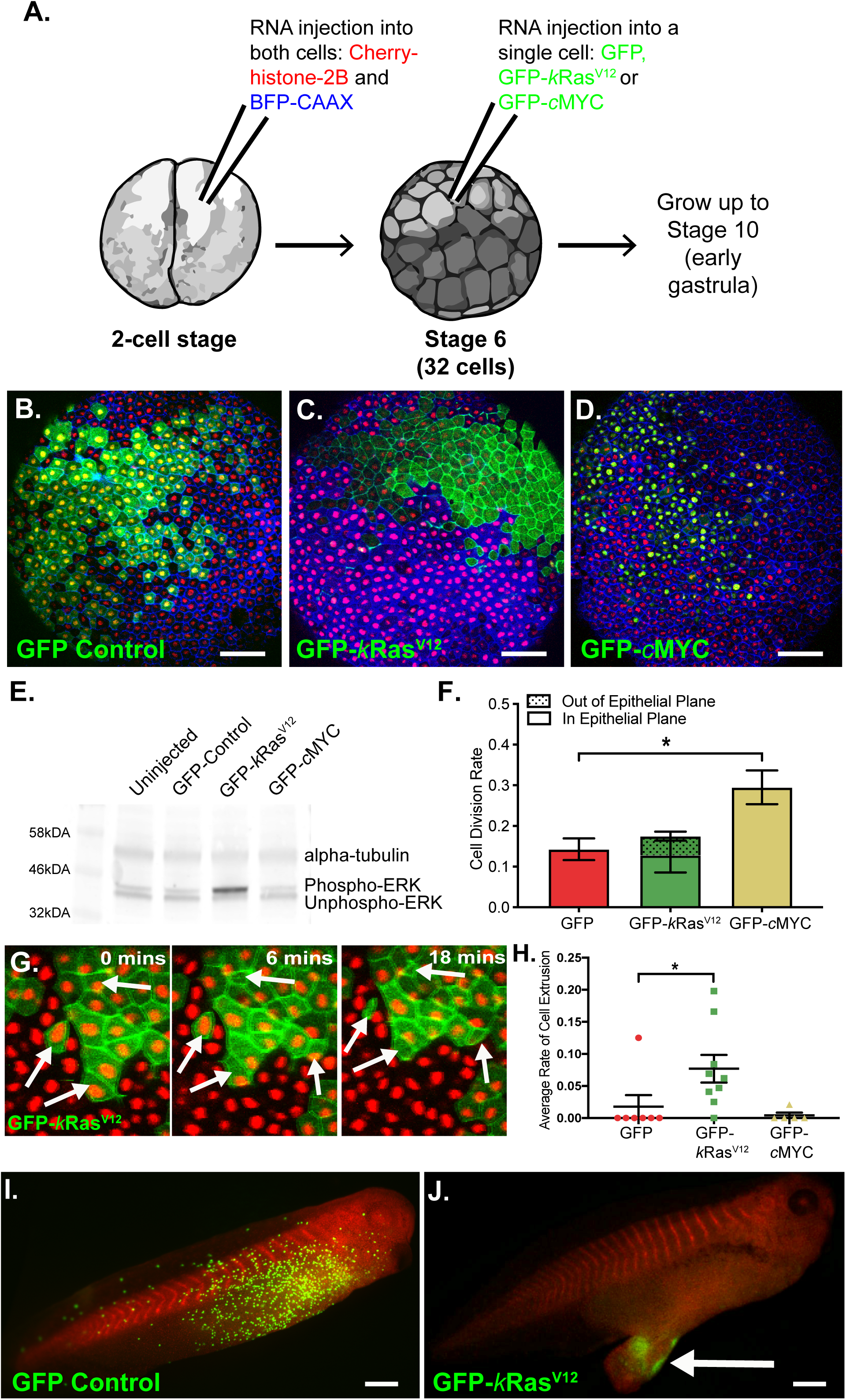
Modelling early stage carcinoma in *Xenopus laevis.* **A.** Schematic of the microinjection protocol. *Xenopus* embryos were injected with Cherry Histone H2B and BFP-CAAX mRNA at the 2-cell stage. At the 32-cell stage, a single cell was injected with GFP, GFP-*k*Ras^V12^ or GFP-*c*MYC mRNA. Embryos were developed to early gastrula stage 10 and imaged. **B-D.** Confocal microscopy images of *Xenopus* embryos developed to early gastrula stage 10, following injection of a single cell at the 32-cell stage with (B) GFP, (C) GFP-*k*Ras^V12^ or (D) GFP-*c*MYC mRNA. **E.** Western blot showing phosphorylated ERK, unphosphorylated ERK and α-tubulin expression in uninjected control embryos and in embryos injected with GFP, GFP-*k*Ras^V12^ or GFP-*c*MYC mRNA at the 32-cell stage. Embryos were lysed at early gastrula stage 10. **F.** Bar chart showing the average percentage of cells that divided per minute of time-lapse, in either GFP, GFP-*k*Ras^V12^ or GFP-*c*MYC overexpression clusters (*p=0.0174 with Kruskal-Wallis test: n=7 GFP-control, 8 GFP-*k*Ras^V12^ and 9 GFP-*c*MYC embryos). Also displayed is the proportion of cell divisions that occurred out of the epithelial plane (shaded portion of the bar). Error bars show SEM. **G.** Stills from a confocal microscopy time-lapse of a representative embryo with a GFP-*k*Ras^V12^ cell cluster at stage 10. White arrows highlight cells observed to be lost basally over the course of the time-lapse. **H.** Dot plot showing average percentage of cells that extruded basally from GFP, GFP-*k*Ras^V12^ or GFP-*c*MYC cell clusters (*p=0.0157 with Kruskal-Wallis test: n=7 GFP, 9 GFP-*k*Ras^V12^ and 5 GFP-*c*MYC embryos). Error bars are SEM. **I-J.** Microscopy images of representative embryos at stage 38 that had a (I) GFP or (J) GFP-*k*Ras^V12^ expressing cluster at stage 10. Anterior is towards the right. Scale bars are 500 μm.

The *k*Ras^V12^ construct was confirmed functional by showing that its expression stimulated an increase in ERK phosphorylation (Figure 1E). As expected from previous studies, *c*MYC significantly increased the cell division rate (CDR) of cells expressing the oncogene, in comparison to control-GFP cells (p=0.0174, Figure 1F). Slightly surprisingly, *k*Ras^V12^ did not increase CDR in expressing cells but, consistent with previous reports from cultured monolayers [14, 15], did increase the propensity for expressing cells to divide out of the epithelial plane (Figure 1F). Contrasting to existing studies [16-19], we did not observe apical extrusion of any *k*Ras^V12^ cells over the course of our time-lapses (Figure S1C) but *k*Ras^V12^ cells were observed being lost basally from the superficial layer (Figure 1G-H). Moreover, imaging of fixed, bisected embryos revealed an increased number of cell layers and cells that had completely delaminated from the tissue (Figure S1D-E). As in previous studies carried out in embryonic contexts, overexpression of *c*MYC did not induce apoptosis alongside the documented increase in cell division (Figure S1F) [20, 21].

When embryos with *k*Ras^V12^ clusters were developed to later stages, the formation of induced tumour-like structures (ITLS) was observed, as described previously [22], whereas embryos with control-GFP clusters were morphologically normal (Figure 1I-J). Embryos with *c*MYC clusters did not develop ITLS, however we observed fast turnover of *c*MYC as almost no GFP-positive cells were retained at these later stages (Figure S1G) [23, 24].

### *k*Ras^V12^ cell clusters have altered mechanical properties and impose a strain on the wild-type epithelium

Having confirmed many of the expected activities of *k*Ras^V12^ and *c*MYC in our oncogene-expressing clusters, we next investigated how the mechanical properties of the tissue might be altered. Previous work has indicated that cells expressing oncogenic Ras are hypercontractile, stiffer than normal cells and exert increased traction forces on their substrate [11-13, 25-27]. We therefore tested how the mechanical properties of the cell cortex in the oncogene-expressing cells were affected in our system by measuring recoil of the junctional vertices following laser ablation of the cell edge (Figure 2A-D). Analysis of vertex-vertex recoil over time allows inferences about the mechanical properties of the cortex to be made [28]. Assuming that the cortex behaves as a viscoelastic Kelvin-Voight material, and that the viscosity of the cytoplasm remains constant (see Material and Methods), the initial recoil velocity reflects the contractile force experienced by the junction prior to ablation. Using laser ablation, we found that the initial recoil of cells in *k*Ras^V12^ clusters was significantly higher when compared to control-GFP or *c*MYC clusters or wild-type cells in tissue surrounding *k*Ras^V12^ clusters (Figure 2A-D), indicating higher cortical contractility in *k*Ras^V12^ cells.

**Figure 2:**
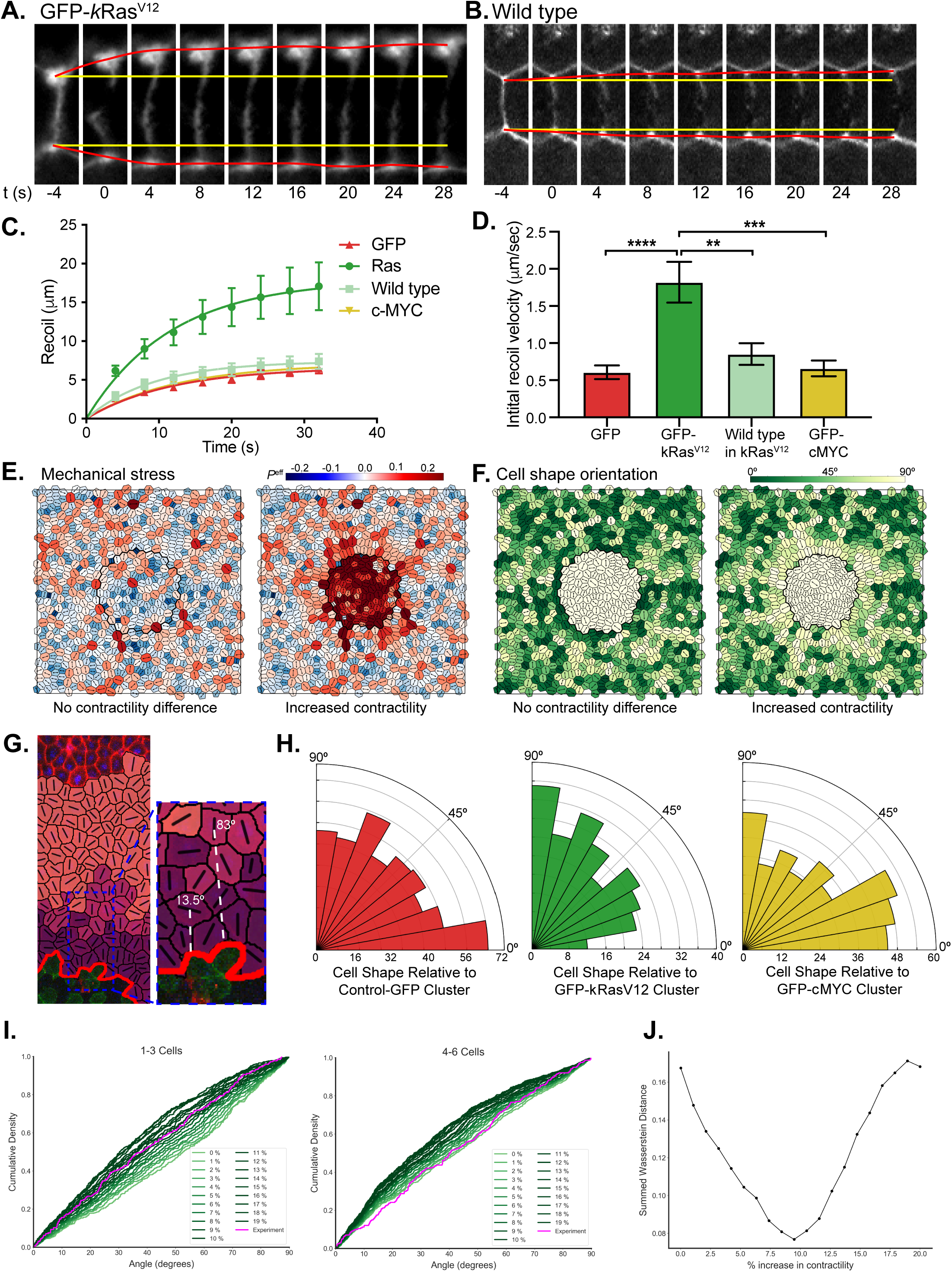
*k*Ras^V12^ cell cluster imposes a mechanical strain on the wild-type epithelium. **A-B.** Cropped regions of confocal movie stills showing laser ablation at a cell edge (highlighted by cherry-UtrCH: F-actin) in a GFP-*k*Ras^V12^ cluster (A) and a surrounding wild-type cell (B). Ablation occurs at t=0; yellow lines show the original positions of cell vertices before laser ablation; red lines show the real-time positions of cell vertices. **C.** Recoil measurements for cells in GFP-control (red), GFP-*k*Ras^V12^ (green) and GFP-*c*MYC (yellow) clusters and areas of wild-type tissue around GFP-*k*Ras^V12^ clusters (Wild-type; light green); n=10 cells for each sample, error bars are SEM. **D.** Initial recoil velocity calculated from recoil measurements in (C); One-way ANOVA: **p<0.01, ***p<0.001, ****p<0.0001, n=10 cells for each sample, error bars are SEM. **E.** Simulated tissue, randomly generated, starting under conditions of zero net tissue stress. Heatmap indicates magnitude of cell-level isotropic stress, *P*^eff^, with cells being under net tension (red) or compression (blue). A simulated Ras cluster was initialised in the centre of the tissue (enclosed within black ring). Left: no additional contractility in cluster; right: 30% increase in cortical contractility, Γ, in cluster. **F.** Simulated tissues from E, with heatmap showing the orientation of the principal axis of cell shape relative to the cluster (as shown in G). **G.** From confocal images, the shapes of host cells neighbouring the clusters (dark purple: 1-3 cells from cluster; light purple 4-6 cells; pink 7+cells) were traced and cell shape orientation (long axis) relative to the cluster was measured (two examples in white are shown). **H.** Rose histograms showing the orientation of wild-type cells’ long-axes 1-3 cells from, GFP-control (red), GFP-*k*Ras^V12^ (green) and GFP-*c*MYC (yellow) clusters, relative to the cluster, with the total number of cells analysed across all embryos in each data group in 10° bins. Kruskal-Wallis test: GFP vs. GFP-*k*Ras^V12^ p=0.0072, GFP vs. GFP-*c*MYC p>0.9999; n=431 cells from 7 GFP embryos, 224 cells from 5 GFP-*k*Ras^V12^ embryos and 348 cells from 7 GFP-*c*MYC embryos. **I.** Cumulative distributions of cell shape orientation relative to cluster (as shown in G), from experiments (magenta) and simulations (green). Ras clusters were simulated with varying degrees of increased cortical contractility, Γ. **J.** Wasserstein distance between experiments and simulations for cumulative distributions in I and S2E. Discrete intervals on the x-axis relate to shades of green in I. For every contractility interval, the y-axis shows the sum of the Wasserstein distances over the three distance categories (1-3, 4-6 and 7+ cells). The best fit is found at a 9% increase in contractility, where summed Wasserstein distance is minimized.

With our recoil measurements indicating that *k*Ras^V12^ cells were significantly more contractile than their surrounding wild-type neighbours, we next investigated how this difference in contractility might affect the distribution of mechanical stress and strain across the tissue. To do this, we adopted a popular vertex-based model of a planar epithelium (see Materials and Methods), to simulate an increase in the cortical contractility of cells in a cluster relative to host tissue. The increased contractility led to a net tensile stress towards the cluster at the interface between neighbouring wild-type cells (Figure 2E), distorting wild-type cell shapes and orienting their long axis towards the cluster (Figure 2F). Importantly, the model predicts that the principal axis of cell shape, as defined by the position of its tricellular junctions (hereto referred to as the long-axis), aligns exactly with the major axis of cell-level stress, for a cell with homogenous and isotropic material properties [29]. This theoretical result has been well-approximated by multiple force inference methods [30-32]. We therefore used the long axis of cell shape as an indicator of the principal axis of mechanical stress.

To compare the model predictions with experimental data, we measured cell shape in neighbouring wild-type cells around GFP, *k*Ras^V12^ and *c*MYC clusters (Figure 2G). As predicted by the model, cells around the *k*Ras^V12^ clusters had altered cell shape, with wild-type cells up to 3 cell-widths from the cluster being significantly more likely to have their long-axes oriented in the direction of the cluster, compared with equivalent cells in control-GFP embryos (p=0.0072, Figure 2H). As discussed above, the model predicts that this is synonymous with a reorientation of cell stress [29]. On the other hand, wild-type cells up to three cells from *c*MYC clusters did not show a significant difference in the orientation of their long-axes (Figure 2H), as would be expected since our recoil experiments found no change in the contractility of *c*MYC cells (Figure 2C-D). The change in wild-type geometry in *k*Ras^V12^ embryos was a localised effect: more than three cells from *k*Ras^V12^ clusters, the orientation of cells’ long-axes was not significantly different to equivalent cells in control-GFP embryos (Figure S2A-D).

To further interrogate the changes in mechanics and shape, we combined our experimental data with the vertex model to predict the contractility change required in the cluster to elicit the cell shape changes seen around the *k*Ras^V12^ clusters. Multiple simulations were run, where the contractility of the *k*Ras^V12^ cluster was simulated by gradually increasing the cortical stiffness parameter from 0% to 20% (see Materials and Methods). Grouping cells into three categories by their distance away from the cluster (in cell widths: 1-3, 4-6 & 7+), we compared cumulative distributions of the orientations of cell shapes, relative to the direction to cluster, between simulations and experiments (Figure 2I and S2E). To determine which simulation best matched the experimental data we calculated the Wasserstein distance (lowest Wasserstein distance is the closest match) between the simulated and experimental cumulative distributions, summed over the three distance categories. We found that a 9% increase of the cortical stiffness parameter in the cluster perturbed local surrounding cell shape orientations to the degree observed in our experimental data (Figure 2J). Our simulations demonstrate that increased contractility can lead to a radial stress that generates an anisotropic strain across the local tissue environment, altering surrounding cell shapes.

Taken together, these data suggest that cells in *k*Ras^V12^ clusters are more contractile than their surrounding neighbours and that these differences lead to cells around the cluster being pulled and distorted in the cluster’s direction. These distortions indicate that the presence of *k*Ras^V12^, but not GFP or *c*MYC, leads to a radial stress across the epithelium which is oriented towards the cluster.

### Wild-type epithelium responds to oncogene-expressing clusters with altered cell division

Cell division is known to be extremely sensitive to tissue tension [1, 33-36] and anisotropic strain in particular: stretching an epithelium increases cell division rate and reorients divisions along the axis of strain [3, 5, 37]. Since the radial stress induced by *k*Ras^V12^ clusters generates anisotropic strain, we hypothesized that this could lead to cell division changes in the host tissue. To test our hypothesis, we used time-lapse confocal microscopy to investigate cell division rate (CDR) and orientation (CDO) in wild-type cells around GFP, *k*Ras^V12^ and *c*MYC clusters. We found that wild-type cells up to three cell widths from *k*Ras^V12^ showed a significant increase in CDR, compared to equivalent cells in control-GFP embryos (p=0.0476, Figure 3A). Somewhat surprisingly, considering we saw no significant effect on cortical contractility (Figure 2C-D) or cell shape (Figure 2H), we found that *c*MYC cell clusters had an even more pronounced effect on CDR in surrounding wild-type cells, with an approximate 4-fold increase in CDR compared to equivalent cells in control-GFP embryos (p=0.0003, Figure 3A). In both *k*Ras^V12^ and cMYC embryos this was not a boundary-specific effect, as the CDR of wild-type cells in direct contact with the oncogene-expressing cells was not significantly different to that two to three cells away (Figure 3B). The induction of division in the wild-type epithelium was a localised effect, with cells more than six cell widths from either *k*Ras^V12^ or GFP-*c*MYC clusters not displaying a significant difference in CDR, compared with control-GFP embryos (Figure 3A).

**Figure 3:**
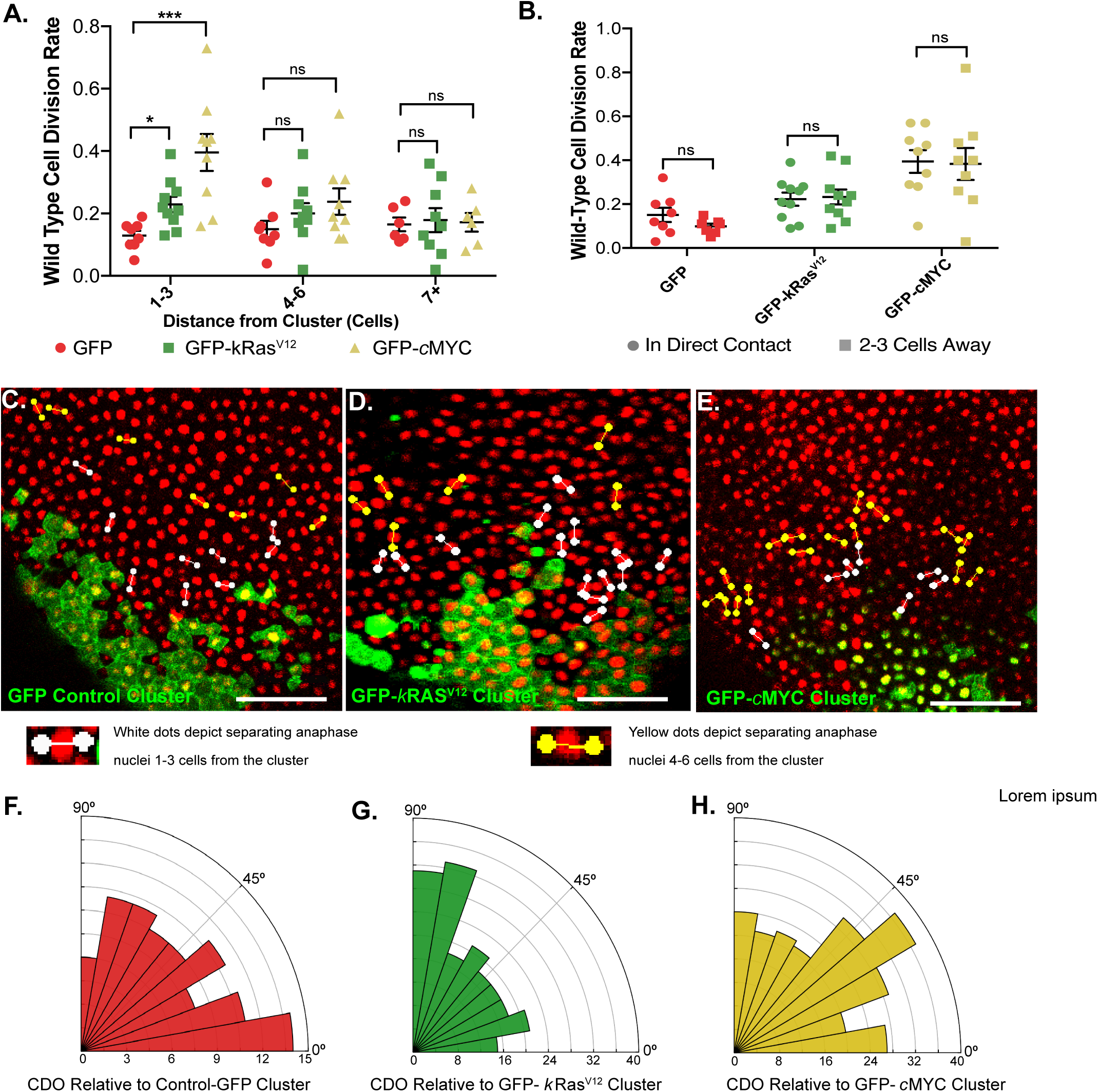
The wild-type epithelium responds to oncogene-expressing clusters with altered cell division. **A-B.** Dot plots showing the percentage of wild-type cells that divided per minute of time-lapse at different distances from GFP, GFP-*k*Ras^V12^ or GFP-*c*MYC clusters. (A) Kruskal-Wallis test: *p=0.0476 and ***p=0.0003, n=8 GFP-control, 10 GFP-kRas^V12^ and 9 GFP-*c*MYC embryos. Error bars are SEM. (B): Paired t-tests were performed, n=8 GFP, 10 GFP-*k*Ras^V12^ and 9 GFP-*c*MYC embryos. **C-E.** Snapshots from confocal microscopy time-lapses of representative embryos showing the orientation of cell divisions that occurred in wild-type cells: coloured lines were drawn connecting the dividing anaphase nuclei. White lines label divisions 1-3 cells from the cluster and yellow lines mark divisions 4-6 cells away. Scale bars are 100 μm. **F-H.** Rose histograms showing cell division orientation relative to (F) GFP control (G) GFP-*k*Ras^V12^ and (H) GFP-*c*MYC clusters, with the total number of cell divisions analysed across all embryos in each data group in 10° bins. Kruskal-Wallis test: GFP vs. GFP-*k*Ras^V12^ p=0.0136, GFP vs. GFP-*c*MYC p>0.9999; n=88 divisions from 8 GFP embryos, 193 divisions from 11 GFP-*k*Ras^V12^ embryos, 231 divisions from 9 GFP-*c*MYC embryos.

The orientation, as well as the frequency, of cell division is usually tightly controlled within epithelial tissues and is vital for maintaining normal tissue architecture [38]. While the *k*Ras^V12^-expressing cells showed an increased propensity to divide out of the epithelial plane (Figure 1F), wild-type cells up to three cells from both *k*Ras^V12^ and *c*MYC clusters retained their strong bias to divide within the plane of the epithelium, with no examples of wild-type cells dividing out of the epithelial plane observed in any of the time-lapses taken (Data not shown, n=10 GFP-*k*Ras^V12^ and 9 GFP-*c*MYC embryos). However, when CDO within the epithelial plane was examined, wild-type cells close to *k*Ras^V12^, but not *c*MYC, clusters exhibited a directional bias towards the oncogenic cluster (Figure 3C-D). We quantified division orientation by measuring the angle between the separating daughter nuclei at anaphase and the line from the cell centroid to the closest cluster edge (Figure S3A). Wild-type cells up to six cells from *k*Ras^V12^ clusters had significantly altered CDO within the epithelial plane, compared with equivalent cells in control-GFP embryos (p=0.0136, Figure 3F-G and Figure S3B). In contrast, wild-type CDO was not significantly altered in cells surrounding *c*MYC clusters (Figure 3H). As with CDR, the *k*Ras^V12^ effect on CDO was local, with wild-type cells more than six cells from the cluster not displaying a significant difference in CDO compared to control-GFP embryos (Figure S3C-D).

Given that cell division was altered in the host epithelium, as well as in the oncogene-expressing cells, we next investigated whether wild-type cells contribute to the ITLS observed in later stage *k*Ras^V12^ embryos. In order to investigate this, cells that neighboured the GFP-*k*Ras^V12^ mRNA injected cell, at the 32-cell stage, were injected with mCherry-H2B mRNA. At early gastrula stage, the embryos were screened and those with a GFP-*k*Ras^V12^ cell cluster in their superficial layer that was surrounded by mCherry-H2B expressing cells, but not expressing mCherry-H2B in the *k*Ras^V12^ cluster itself were selected (Figure S3E). At stage 38, the *k*Ras^V12^ driven ITLS were analysed and cells expressing mCherry-H2B were found localised to the growths, demonstrating that cells derived from the host epithelium, as well as cells that expressed *k*Ras^V12^, contributed to the tumour-like phenotype (Figure S3F).

These results show that the host epithelium displays altered cell division in response to groups of cells that overexpress *k*Ras^V12^ or *c*MYC. In the case of *k*Ras^V12^, the division effects were similar to that seen when an anisotropic strain is applied to an epithelial tissue: increased division rate and divisions oriented along the principal axis of shape [3]. Moreover, the local nature of these division effects mirrored the changes seen in cell shape around the Ras^V12^ clusters. In contrast, *c*MYC clusters appeared to elicit only an increase in division rate, without perturbing cell shape or division orientation.

### Activation of RhoA in a cell cluster induces a response in the wild-type epithelium comparable to *k*Ras^V12^ expression

Anisotropic stress and strain can be generated when neighbouring tissues with higher levels of actomyosin contractility exert pulling forces [1, 39-42]. Previous reports have shown that expression of oncogenic Ras stimulates non-transformed mammary epithelial cells to exert increased traction forces on their substrate, in a manner dependent on the activity of Rho and non-muscle myosin ll [13]. Myosin ll generates contractile forces by crosslinking actin filaments into higher-order structures and hydrolysing ATP to pull on the fibres [43-47] and Ras-expressing cells exhibit increased phosphorylation of myosin ll in numerous contexts [10, 13, 16]. Fittingly, we found increased phosphorylated myosin ll at tricellular vertices in the *k*Ras^V12^-expressing cells, compared with wild-type cells in these same embryos (Figure 4A). Furthermore, in the *k*Ras^V12^ clusters, F-actin organisation was less homogenous, with an increase in cortical actin close to tricellular vertices (Figure 4B). Importantly, this actin stain did not reveal evidence of an actin ‘purse string’, nor the formation of lamellipodia and filopodia in the wild-type epithelium close to the *k*Ras^V12^ cell cluster, making it unlikely that the change in wild-type epithelial cell shape occurred as the result of a wound-healing-like response [40, 48-50].

**Figure 4:**
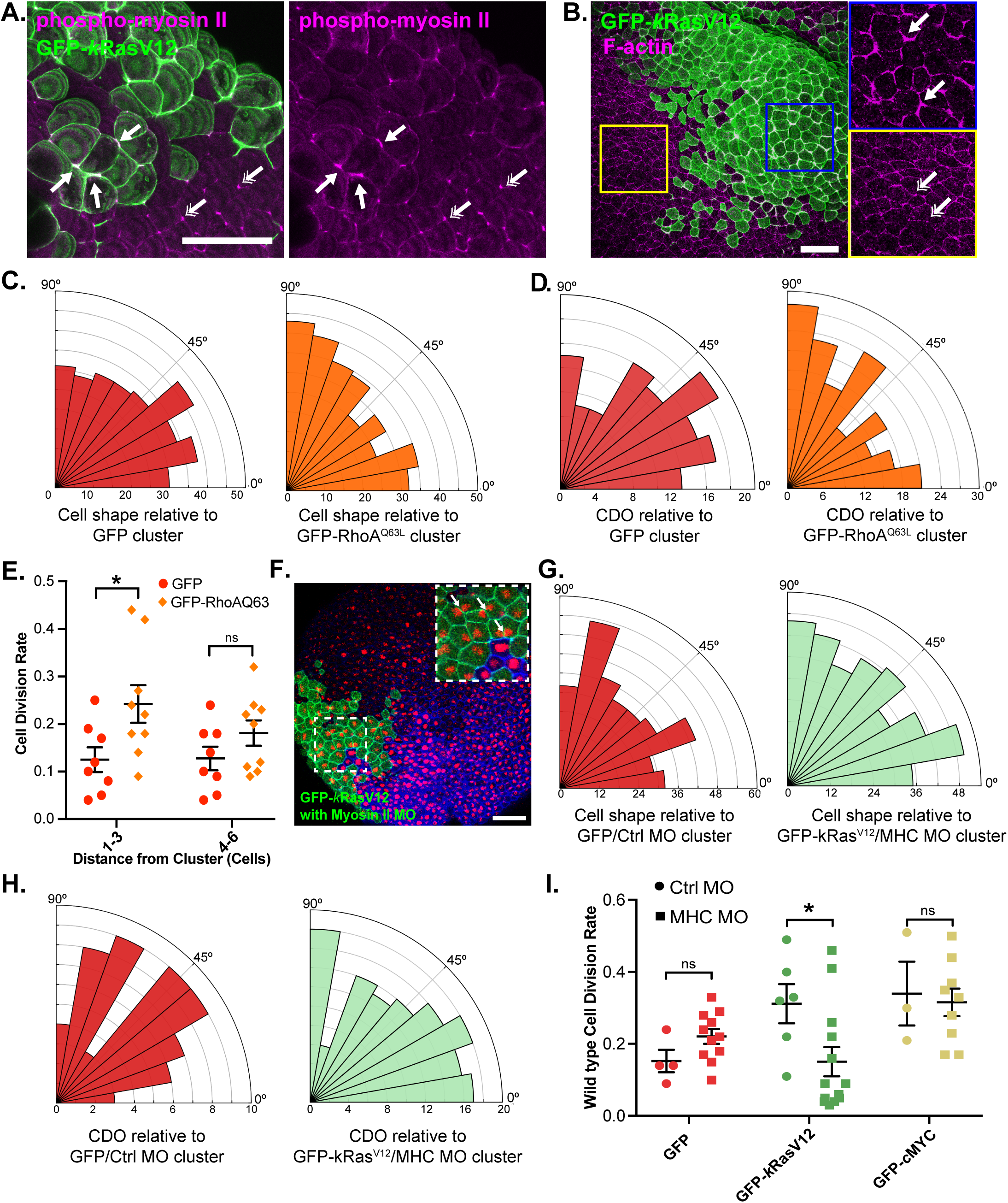
Actomyosin contraction in cluster is required to generate strain and alter cell division in wild-type tissue. **A and B.** Confocal images of fixed, stage 10 embryos with a GFP-*k*Ras^V12^ cluster, stained for **A.** Phosphorylated myosin ll (magenta), single-headed arrows highlight tricellular junctions with increased phospho-myosin ll in GFP-*k*Ras^V12^ cells compared to wild-type tissue (double-headed arrows); **B.** F-actin (phalloidin; magenta), single-headed arrows highlight increased F-actin at the cell cortex in the GFP-*k*Ras^V12^ cluster compared to wild-type tissue (double-headed arrows). **C** Rose histograms showing the orientation of wild-type cells’ long-axes up to 6 cells from GFP-control (red) or GFP-RhoA^Q63L^ (orange) cell clusters, relative to the cluster, with the total number of cells that were analysed across all embryos in each data group in 10° bins. Kolmogorov-Smirnov test: p=0.0468, n=298 cells from 6 GFP-control embryos and 299 cells from 6 GFP-RhoA^Q63L^ embryos. **D.** Rose histograms showing cell division orientation relative to GFP-control (red) or GFP-RhoA^Q63L^ (orange) clusters, with the total number of cell divisions that were analysed across all embryos in each data group in 10° bins. Kolmogorov-Smirnov test: p=0.0368, n=98 divisions from 10 GFP-control embryos and 174 divisions from 9 GFP-RhoA^Q63L^ embryos. **E.** Dot plot showing percentage of wild-type cells that divided per minute of time-lapse at different distances from GFP-control or GFP-RhoA^Q63L^ clusters. One-way ANOVA: *p=0.0250, n=7 GFP-control and 9 GFP-RhoA^Q63L^ embryos. Error bars are SEM. **F.** Confocal microscopy image shows a myosin ll deficient GFP-*k*Ras^V12^ cell cluster. Arrows highlight ‘butterfly-nuclei’. **G**. Rose histograms showing the orientation of wild-type cell long-axes up to 3 cells from GFP/Ctrl MO (red) or myosin ll deficient (MHC MO) GFP-*k*Ras^V12^ (light green) cell clusters, with the total number of cells analysed grouped in 10° bins. Kruskal-Wallis test: p>0.9999, n=325 cells from 6 GFP/Ctrl MO embryos and 368 cells from 7 GFP-*k*Ras^V12^/MHC MO embryos. **H.** Rose histograms show cell division orientation up to 6 cells from (D) GFP/Ctrl MO (red) or GFP-*k*Ras^V12^/MHC MO (light green) cell clusters, with the total number of cell divisions that were analysed across all embryos in each data group in 10° bins. Kruskal-Wallis Test: p=0.9327, n=58 divisions from 6 GFP/Ctrl MO embryos and 132 divisions from 9 GFP-*k*Ras^V12^/MHC MO embryos. **I.** Dot plot shows percentage of wild-type cells that divided per minute of time-lapse, up to 3 cells from GFP, GFP-*k*Ras^V12^ or GFP-*c*MYC control morpholino clusters, or myosin ll deficient GFP, GFP-*k*Ras^V12^ or GFP-*c*MYC clusters. Kruskal-Wallis test: *p=0.0319, n=5 GFP/Ctrl MO embryos, 11 GFP/MHC MO, 6 GFP-*k*Ras^V12^/Ctrl MO, 13 GFP-*k*Ras^V12^/MHC MO, 3 GFP-*c*MYC/Ctrl MO and 9 GFP-*c*MYC/MHC MO embryos. Scale bars in A, B and F represent 100 µm.

Myosin ll is phosphorylated downstream of RhoA activation, which occurs in response to constitutive activation of Ras [51-55]. To examine whether activation of RhoA is sufficient to induce anisotropic strain in surrounding wild-type tissue, a group of cells were generated expressing the constitutively active RhoA Q63L mutant [56, 57]. Wild-type cells up to three cells from RhoA^Q63L^ clusters were more likely to have their long-axes oriented in the cluster’s direction, compared with analogous cells in control-GFP embryos (p=0.0468, Figure 4C). Furthermore, the CDO of wild-type cells up to six cells from RhoA^Q63L^ clusters was altered, with cells showing an increased propensity to orient divisions towards the cluster (p=0.0368, Figure 4D). CDR was also significantly increased in wild-type cells up three cells from RhoA^Q63L^ clusters, (p=0.0250, Figure 4E). These results demonstrate that a group of cells with increased RhoA activity is sufficient to induce cell shape changes in the surrounding wild-type epithelium, indicative of anisotropic strain. Furthermore, the presence of a group of cells with increased RhoA activity can alter wild-type cell division in a similar manner to a *k*Ras^V12^ cluster – increasing the CDR and inducing directional bias to division orientation within the epithelial plane.

### Non-muscle myosin ll is required in *k*Ras^V12^-expressing cells for the cluster to alter wild-type tissue mechanics and cell division

Non-muscle myosin ll is required for epithelial cells to generate contractile forces [43-47]. To directly test whether *k*Ras^V12^ cell contractility induces the observed anisotropic strain in the surrounding wild-type epithelium, we specifically knocked down myosin ll in the *k*Ras^V12^-expressing cells, through injection of a morpholino (MHC MO) in these cells only [47]. The presence of butterfly-shaped nuclei in the *k*Ras^V12^ clusters indicated that myosin ll levels were reduced, with cells struggling to complete cytokinesis due to decreased functionality of the actomyosin contractile ring (Figure 4F). The myosin ll knockdown did not significantly affect the cell-autonomous CDR of *k*Ras^V12^ cells (Figure S4A), despite the length of mitosis being significantly longer (p<0.0001, Figure S4B). Crucially, when myosin ll was knocked-down in the *k*Ras^V12^ cells, we found that wild-type cells up to three cells from these myosin ll-deficient *k*Ras^V12^ clusters had their long-axes oriented uniformly, with the distribution of long axis orientation no longer significantly different to equivalent cells in control embryos (Figure 4G). Therefore, knockdown of myosin ll in the *k*Ras^V12^ cells recovered cell shape in the surrounding wild-type epithelium, indicating that isotropic stress had been restored at the tissue level. Knockdown of myosin ll in GFP or *c*MYC clusters had no effect on cell shape: surrounding wild-type cells still displayed uniformly oriented long-axes, with no significant difference to control embryos (Figure S4C-D).

Given that surrounding wild-type cell geometry was rescued, we investigated whether cell division was also recovered. When myosin ll was knocked down in the *k*Ras^V12^ cells, wild-type cells up to six cells from these clusters lost the biased CDO seen around *k*Ras^V12^ clusters and instead divided uniformly in all directions, no longer significantly different to control GFP clusters (p=0.4013, Figure 4H). Myosin ll knockdown had no effect on wild-type CDO in GFP or *c*MYC clusters, with cell divisions uniformly oriented within the epithelial plane (Figure S4E-F). Furthermore, the CDR of wild-type cells close to myosin ll-deficient *k*Ras^V12^ clusters was significantly reduced compared to wild-type cells close to control-MO *k*Ras^V12^ clusters (p=0.0299, Figure 4I). In contrast, knockdown of myosin ll in *c*MYC clusters did not significantly affect surrounding wild-type CDR (p=0.7908, Figure 4I). Together these data indicate that myosin ll is required in *k*Ras^V12^ cells in order for a *k*Ras^V12^ cluster to generate anisotropic strain in the surrounding host epithelium and lead to increased CDR and altered CDO in these wild-type cells.

In conclusion, we find that the presence of *k*Ras^V12^ clusters in an otherwise normal epithelium leads to the generation of localised anisotropic strain oriented towards the cluster, as observed by changes in cell shape orientation around the cluster. Anisotropic strain is known to alter cell division dynamics [3, 5, 37] and, consistent with this, we see increased cell division rate and altered division orientation in wild-type cells around *k*Ras^V12^ clusters. We find that the induced anisotropic strain is caused by a radial tension, produced by the difference in actomyosin contractility between *k*Ras^V12^ cells and the surrounding host epithelium. Isotropy and normal cell division dynamics can be recovered in the wild-type tissue by reducing contractility specifically in the *k*Ras^V12^ cells by knockdown of myosin II.

In addition to changes in cell division around *k*Ras^V12^ clusters, we also find that host cell division rate is significantly increased around *c*MYC clusters, while division orientation and cell shape remained unaffected. Since knockdown of myosin ll in the *c*MYC cells did not recover surrounding wild-type cell division rate, we suggest that a distinct mechanism drives increased host cell division with *c*MYC compared to *k*Ras^V12^. Since previous studies have shown that *c*MYC-overexpression alters a cell’s secretome and inhibits the secretion of anti-mitotic factors [58, 59], a likely possibility is that the host cells around the *c*MYC clusters are responding to a change in their chemical, rather than mechanical, environment.

Together, these results indicate novel roles for *k*Ras and *c*MYC in inducing and dysregulating cell division in a host epithelium. An exciting avenue for future research is to determine whether the same responses occur in differentiated, adult tissues during carcinoma onset. The dysregulation of wild-type cell division in host epithelia could help drive the increase in cell number that defines early cancer stages and aid the spread of oncogenic cells through epithelial crowding and cell delamination [2, 60]. Moreover, faster and dysregulated divisions in the host tissue could also increase the chance of these cells acquiring genetic changes of their own. This would imply that primary tumour development, and even the onset of secondary metastases, could be multi-focal, arising from genetic damage in multiple cells. As tumour heterogeneity is central to drug resistance [61], targeting this co-opting of the host epithelium could help to make therapeutic interventions more effective.

## MATERIALS AND METHODS

### Constructs

Human *k*RasV12 and *c*MYC were used in these experiments (Table 1). *k*Ras is 82% conserved at the mRNA level between *Xenopus* and mammals, with the proteins encoded sharing highly similar structures [62]. *c*MYC is also highly conserved across vertebrates, including *Xenopus* [63] and human *c*MYC has previously been demonstrated to rescue phenotypes induced in *Xenopus* when endogenous *c*MYC function is abrogated [64]. Both constructs are also fusion proteins, N-terminally tagged with GFP (Table 1). *k*Ras had been N-terminally tagged in numerous studies, with no apparent consequences on its functionality [65]. *c*MYC has also been N-terminally tagged with GFP in numerous studies, with one study showing GFP-*c*MYC can functionally replace endogenous *c*MYC in mice [66].

**TABLE 1:** Plasmids List.

| Plasmid | Sources |
| --- | --- |
| pCS2 Cherry-Histone 2B | Woolner Lab Stocks |
| pCS2 BFP-CAAX | Gift from the Bement Lab [67] |
| pCS2 N-GFP | Woolner Lab Stocks |
| pCS2 GFP-*kRas*<sup>V12</sup> | pBabe human *kRas*<sup>V12</sup> was purchased from Addgene #2544 and cloned into pCS2 N-GFP (above). PCR primers were used to add 5' BspE1 site and 3' STOP codon and Xho1 site. |
| pCS2 GFP-*cMYC* | MSCV human *cMYC*-IRES-GFP was purchased from Addgene #8119 and cloned into pCS2 N-GFP (above). PCR primers were used to add 5' BspE1 site and 3' STOP codon and Xho1 site. |
| pcDNA3-EGFP-RhoA-Q63L | Purchased from Addgene #12968 |

### mRNA Synthesis

Plasmids were linearised by restriction enzyme digestion. The resultant linearised DNA was the purified by a phenol/chloroform extraction and *in vitro* capped mRNA synthesis was carried out according to manufacturer’s instructions *(Ambion, #AM1340).* mRNA was then purified by a phenol/chloroform extraction. mRNA was diluted to 1 μg/μl and stored at −80°C until use.

### Priming *Xenopus laevis*

Female *Xenopus laevis* were pre-primed with 50 units of Pregnant Mare’s Serum Gonadotrophin (Intervet UK) injected into the dorsal lymph sac. Four to seven days later, frogs were then primed with 500 units of Human Chorionic Gonadotrophin (Intervet UK) injected into the dorsal lymph sac [68]. Each frog was housed individually overnight and approximately 18 hrs later, the *Xenopus laevis* were transferred into room temperature (RT) ‘high salt’ 1x Marc’s Modified Ringers (MMR) solution (100 mM NaCl, 2 mM KCl, 1 mM MgCl, and 5 mM HEPES [pH 7.4]). Eggs were collected from tanks 2-5 hours later.

### Embryo Fertilisation

*In vitro* fertilisation was performed as described previously [68]. MMR was removed from the collected eggs. A small amount of testis prep was cut up and spread over collected eggs to ensure all were exposed. After 5 mins at RT, the dish was topped up with 0.1X MMR and left for a further 30 mins. MMR was then drained and the embryos transferred into a glass beaker. 50 ml of 2% L-cysteine solution (2 g L-cysteine *(Sigma Aldrich, #168149-100G)* in 100 ml 0.1% MMR, pH 7.8 – 8.0) was added, and swirled gently until the jelly coat of the embryos was reduced. The L-cysteine solution was removed and the embryos washed a minimum of six times, with a total 200 ml 0.1% MMR. The embryos were transferred into new 10 ml petri dish and topped up with fresh 0.1% MMR then incubated at RT to reach 2-cell stage.

### mRNA Microinjection

Microinjections were carried out using Picospritzer lll Intracel injector (Parker instrumentation). Healthy embryos at the 2-cell stage were transferred into an injection dish containing 0.1X MMR with 5% Ficoll *(SigmaAldrich, #PM400)*. Each cell was injected with a total volume of 4.2 nl (for constructs and concentrations injected, see Table 2 below). Following this microinjection, embryos were washed in a Petri dish containing 0.1% MMR, then transferred into a second Petri dish containing fresh 0.1% MMR. These embryos were left at RT to develop to the 32-cell stage. At the 32-cell stage, the embryos were transferred back into the injection dish, containing 0.1% MMR and 0.5% Ficoll, and cells at the animal pole were injected with a total volume of 2.1 nl (for constructs and concentrations injected, see Table 2 below). Following microinjection, the embryos were washed in a Petri dish containing 0.1% MMR and then transferred into a second Petri dish containing fresh 0.1% MMR and incubated at 16°C overnight.

**TABLE 2:**
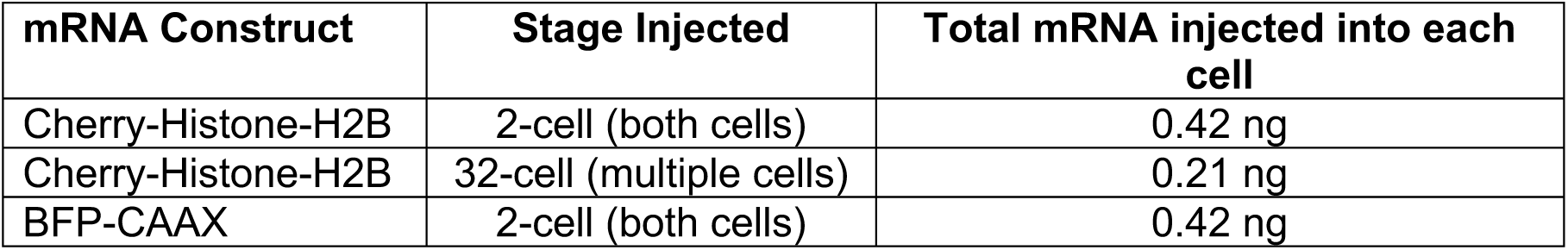
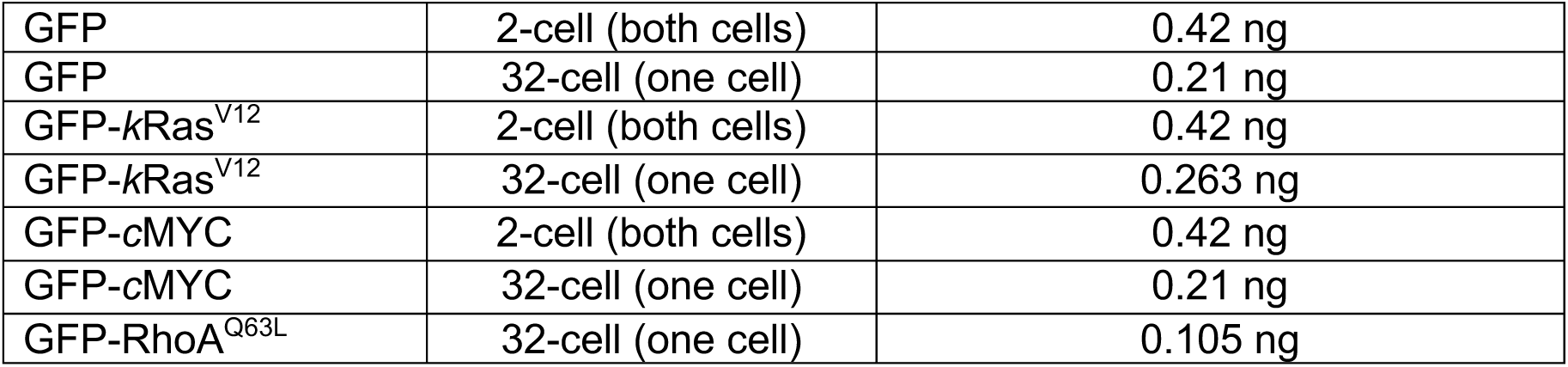
List of mRNA concentrations injected into *Xenopus* embryos.

### Myosin ll Knockdown

Myosin ll was knocked down through microinjection of a Morpholino targeting non-muscle myosin ll heavy chain 2B (MHC) (Table 3) [47]. Prior to microinjection, the Morpholino was heated for 10 minutes at 65°C and combined with GFP-*k*Ras^V12^ mRNA. The final needle concentration of the morpholino was 0.2 μM. A single cell at the animal pole was injected with a total volume of 2.1 nl of the GFP-Ras^V12^ mRNA (above) and MHC Morpholino.

**TABLE 3:**
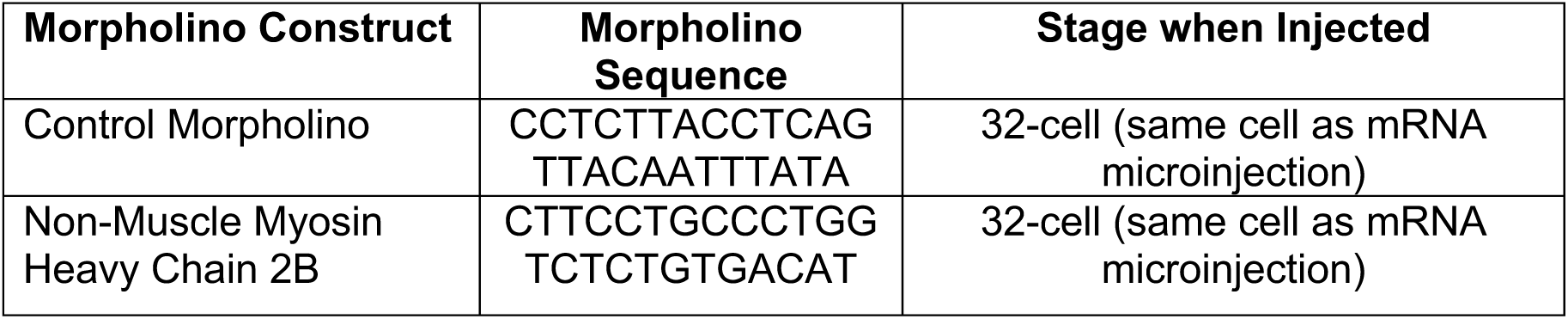
List of morpholinos injected into *Xenopus* embryos.

### Embryo Survival and Cluster Quantification

Following microinjection, embryos were left at 16°C for 16 hrs. At stage 10 [69] embryos were screened for survival and for the presence of an apical GFP cluster.

### Western Blotting

Injected embryos were washed three times in PBS and then lysed by pipetting up and down in 10 µl ice-cold lysis buffer (Tris-HCl pH7.5, 150 mM NaCl, 0.5% NP-40, 5 mM EGTA, 5 mM EDTA) supplemented with 1X Protease inhibitor cocktail *(Promega G6521)* and PhosSTOP phosphatase inhibitor *(Sigma Aldrich)* per embryo. The embryos were then spun at 16873 x g for ten mins at 4°C and the supernatant transferred into fresh tubes. Up to 10 µl of each sample was diluted with lysis buffer to make total volume of 15 µl. 5 µl of 4X loading buffer (8% SDS, 0.2 M tris-Cl pH 6.8, 8% Glycerol and 0.8% 2-mercaptoethanol) was added and the samples were incubated at 95°C for five mins. Samples were loaded into 4–20% Mini-PROTEAN TGX Stain-Free Protein Gels *(Bio-Ra, #4568093)* and were fractionated by SDS-PAGE, before transfer to a 0.45 µm nitrocellulose membrane *(GE Healthcare, #10600002)* using a transfer apparatus according to the manufacturer’s protocols *(Bio-Rad)*. The membrane was blocked by incubation with 5% non-fat milk (or 5% BSA for phospho-specific antibodies) in TBST (10 mM Tris, pH 8.0, 150 mM NaCl, 0.5% Tween 20) for 1 hr. Following this, the membrane was washed once with TBST and incubated with primary antibodies at 4°C for 12 hrs (Phospho-ERK1/2 1:500 *(Sigma Aldrich, #E7028)*; ERK1/2 1:1000 *(Cell Signaling, #9102S)*, α-tubulin *(Sigma Aldrich, #T9026)*). The antibodies were diluted in the same solution that was used for blocking. Membranes were washed three times for 10 mins with TBST and incubated with IRDye conjugated antibodies (Goat anti-Rabbit IRDye800CW 1:5000 *(abcam, #216773)*, donkey anti-mouse IRDye680RD 1:5000 *(abcam, #216778)*), diluted in blocking solution. Membranes were then washed three times more and an Odyssey CLX LICOR was used to image the blot.

### Immunofluorescence

Embryos at stage 10 were fixed overnight with a gentle rotation at RT in ‘microtubule fix’, consisting of 3.7% fresh formaldehyde, 0.25% glutaraldehyde, 0.2% Triton X-100, 69.6 mM K-Pipes, 4.35 mM EGTA, 0.87 mM MgCl_2_. The following day, embryos were washed five times with PBS and the vitelline membranes were removed using forceps. Embryos were then quenched in 100 mM sodium borohydride in PBS for 2 hrs, rotating at RT. Embryos were washed three times in PBS for five mins and bleached for 90 mins in 10% H2O2 on a lightbox at RT. Embryos were washed three times for ten minutes on a rotator in TBSN (Tris-buffered saline: 155 mM NaCl, 10 mM Tris-Cl [pH 7.4]; 0.1% Nonidet P-40) and then blocked overnight in 10 mg/ml BSA at 4°C with rotation. The block solution was changed twice the following day, and then primary antibodies were added to the embryos at a dilution of 1:200 (GFP-tag *(Invitrogen, MA5-15256)*, phospho-myosin light chain 2 (S19) 1:500 *(Cell Signalling, #3671 lot 3 and lot 6)*) and incubated overnight at 4°C. The following day, embryos were washed five times in TBSN/BSA for one hr at 4°C whilst rotating and incubated with secondary antibodies overnight at 4°C at a dilution of 1:400 (Alexa Fluor 488 goat anti-mouse *(Invitrogen, #A11001)* and 1:400 Poly-HRP-conjugated secondary antibody from Tyramide SuperBoost Kits with Alexa Fluor 568 (*ThermoScientific, #B40956*)). Embryos were then washed three times in TBSN/BSA for one hr at 4°C whilst rotating and then twice in TBSN alone for one hour at 4°C. Phospho-myosin light chain 2 was visualised using a Tyramide SuperBoost Kits with Alexa Fluor 568 (*ThermoScientific, #B40956*), according to manufacturers instructions. Nuclei were visualised by staining with DAPI at a dilution of 10 µg/ml (*Thermo-Scientific, #D1306*) and then washed three times in TBSN for half an hour at 4°C.

Phalloidin staining was carried out using albino embryos. Injected embryos were rinsed three times in PBS, and then fixed for four hrs at RT (3.7% formaldehyde, 0.25% glutaraldehyde and 0.1% Triton-X in PBS) whilst rotating gently. Embryos were washed three times in PBS and bisected along the sagittal axis using a razor blade and the vitelline membranes removed using forceps. The embryos were washed a further three times in PBTw (PBS + 0.1% Tween) and incubated overnight whilst rotating at 4°C in 0.005 U/µl Alexa Fluor 594 phalloidin (*Invitrogen, #A12381)* in PBTw. The following day, embryos were washed five times in PBS for one hour whilst rotating at 4°C and imaged.

### Live Imaging

Approximately 21 hours after fertilisation, when the embryos were at stage 10 [69], they were transferred into fresh dish of 0.1 X MMR, which had 1 mm Polypropylene mesh (*SpectrumLabs, P/N146410*) stuck to its base to prevent the embryos rolling. Live-imageing was then performed using a dipping lens so as not to apply any mechanical stress by using a coverslip. Images were collected on a Leica TCS SP5 AOBS upright confocal using a 20x/0.50 HCX Apo U-V-I (W (Dipping Lens)) objective and 1x confocal zoom. The confocal settings were as follows: pinhole 1 airy unit, scan speed 1000Hz bi-directional, format 512 × 512. Images were collected using the following detection mirror settings: BFP 406-483 eGFP 498-584 nm and mCherry 604-774 nm using the 405 nm, 488nm (25%) and 594nm (25%) laser lines respectively. Images were collected sequentially to eliminate bleed-through between channels. The distance between each optical stack was maintained at 4.99 μm and the time interval between each capture was 1 min, with each sample imaged for up to 1 hr. The maximum intensity projections of these three-dimensional stacks are shown in the results.

### Laser ablation and recoil measurements

Images of *Xenopus laevis* embryos were acquired using a Nikon A1R confocal microscope using a 60x NA1.4-CFI-Plan-Apo oil objective and NIS-Elements software (Nikon). Laser ablation was performed using a Micropoint ablation laser (Andor Systems) attached to the Nikon confocal. The confocal settings were as follows: pinhole 1 airy unit, scan speed 400Hz unidirectional, format 512 × 512, 1x confocal zoom. GFP and mCherry-UtrCH were imaged using the 488nm and 561nm laser lines respectively. A single focal plane was captured for each embryo with a frame every 4 seconds for 2-3 minutes. A wounding laser level of 85 with a single blast setting was used to create a small wound at the cell edge. In order to provide a pre-ablation image and to visualise the moment of wounding, ablation was performed a few frames into the capture. A cell edge was targeted either within the cluster (at least 3 cells from the perimeter) or within the non-cluster region (at least 3 cells outside of the cluster). To determine initial recoil from the laser ablation movies we followed a previously described protocol [28]. In brief, to measure the deformation of the cell junction following ablation, the xy coordinates of the two vertices (identified by mCherry-UtrCH) each side of the wound were tracked in ImageJ using the MTrackJ plugin. This data was used to extract the initial recoil and *k* (a ratio between junctional elasticity and viscosity of the cytoplasm) values using a Kelvin-Voigt model [70]. No significant difference in *k* values was seen between any of the samples tested (data not shown), meaning that changes in initial recoil could be interpreted as a strong indication that junctional tension was affected [28]. All initial recoil measurements were found to be normally distributed by Shapiro-Wilk test and were compared using a one-way ANOVA with Tukey’s multiple comparisons test.

### Cell Division Analysis

Embryo time-lapse movies were generated using ImageJ64, from which snapshots were selected. Cell division rate in the epithelial plane was quantified as the percentage of cells where daughter nuclei were observed to separate, per minute. Cells that exhibited nuclear envelope breakdown, but where daughter nuclei were not observed to separate within the plane of the epithelium, were assumed to have divided out of plane. In plane CDO was measured using the Image J straight-line tool to draw a line between the dividing nuclei of a cell in anaphase and the closest edge of the cluster. Mitotic length was defined as the time between nuclear envelope breakdown and the first frame where daughter nuclei were observed to separate.

### Cell Shape Analysis

Analysis of cell shapes was carried out by segmenting cells of interest, using an initial manual trace of cell edges. The principal axis of cell shape (described below) was calculated using a previously published in-house Python script [3, 29]. Cell shape was characterised by a shape tensor derived from the second moments of the positions of the tricellular junctions (we also include the rare case where more than three edges meet). For every cell we label the cell vertices *i* = 1,2, …, *n* anticlockwise, where *n* is the number of vertices. The cell centroid, ***C***, is the arithmetic mean of the positions of the tricellular junctions

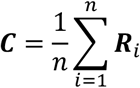

Where ***R***_*i*_ is the position vector of junction *i*. The cell shape tensor, *S*, is then defined as

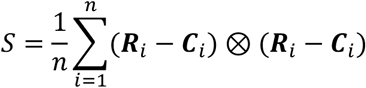

Where ⊗ is the outer product. The principal axis of cell shape is defined as the eigenvector associated with the principal eigenvalue of *S*.

### Simulations using a vertex-based model

Simulations were done in the framework of a vertex-based model, where the tissue is represented as a planar network of polygons. The model and simulation procedure are identical to our previously published methods [3, 29, 71]. Briefly, we assume that every cell has a dimensionless mechanical energy, *U*, defined by

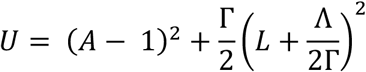

where while *A* and *L* denote the dimensionless area and perimeter of a cell, Γ represents the stiffness of the cortex relative to the bulk and Λ is a mechanical parameter prescribing the preferred perimeter *L*_0_ = −Λ/(2Γ). Mechanical equilibrium is found by minimising the total mechanical energy, summed over all cells. For all simulations we use the parameters (Λ, Γ) = (0.259, 0.172), which have previously been fitted to the Xenopus animal cap tissue [29]. We simulate the effect of extra contractility in Ras clusters by inducing a percentage increase to the reference cortical stiffness parameter, Γ. Such a procedure has previously been shown to well-replicate the behaviour of hyper-contractile tissues [72].

As described in [29], the magnitude of cell stress can be characterised by the isotropic component, *P*^eff^, of the cell-level stress tensor:

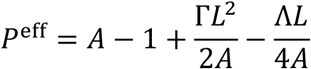

where positive values of *P*^eff^ indicate that the cell is under net tension and negative values indicate net compression.

### Graphs and Statistical analysis

Rose histograms were generated using a python script and all other charts were produced using Prism 7/8 (GraphPad Software, LLC). Statistical analysis was performed using Prism 8. For cell division rate analysis, distributions were first tested for normality using Shapiro-Wilk tests. If normality was passed, a one-way ANOVA with Sidak’s multiple comparisons test was performed; if normality could not be assumed, a Kruskal-Wallis with Dunn’s multiple comparisons test was performed. For comparisons between paired division rate data at the boundary of the oncogenic cluster, distributions were found to be normally distributed by a Shapiro-Wilk test, and paired t-tests were performed. For cell shape and division orientation angle data, normality could not be assumed and individual comparisons were made using Kolmonov-Smirnov tests. For multiple comparisons of angle data, Kruskal-Wallis with Dunn’s tests were performed. For all statistical tests performed, n numbers and p values are given in the relevant figure legends.

## Acknowledgements

MM was supported by a WT 4 Year PhD Studentship [106506/Z/14/Z], ANB was supported by a BBSRC studentship and SW and GKG were supported by a Wellcome Trust/Royal Society Sir Henry Dale Fellowship [098390/Z/12/Z]. The Bioimaging Facility microscopes used in this study were purchased with grants from BBSRC, Wellcome and the University of Manchester Strategic Fund. Thanks to Peter March and Roger Meadows for their help with microscopy and William Bement (University of Wisconsin-Madison) for his kind gift of the BFP-CAAX construct. Also, special thanks to Angeliki Malliri, Paul Martin and Andrew Gilmore for their critical reading of the manuscript.

## SUPPLEMENTARY FIGURE LEGENDS

**Figure S1:**
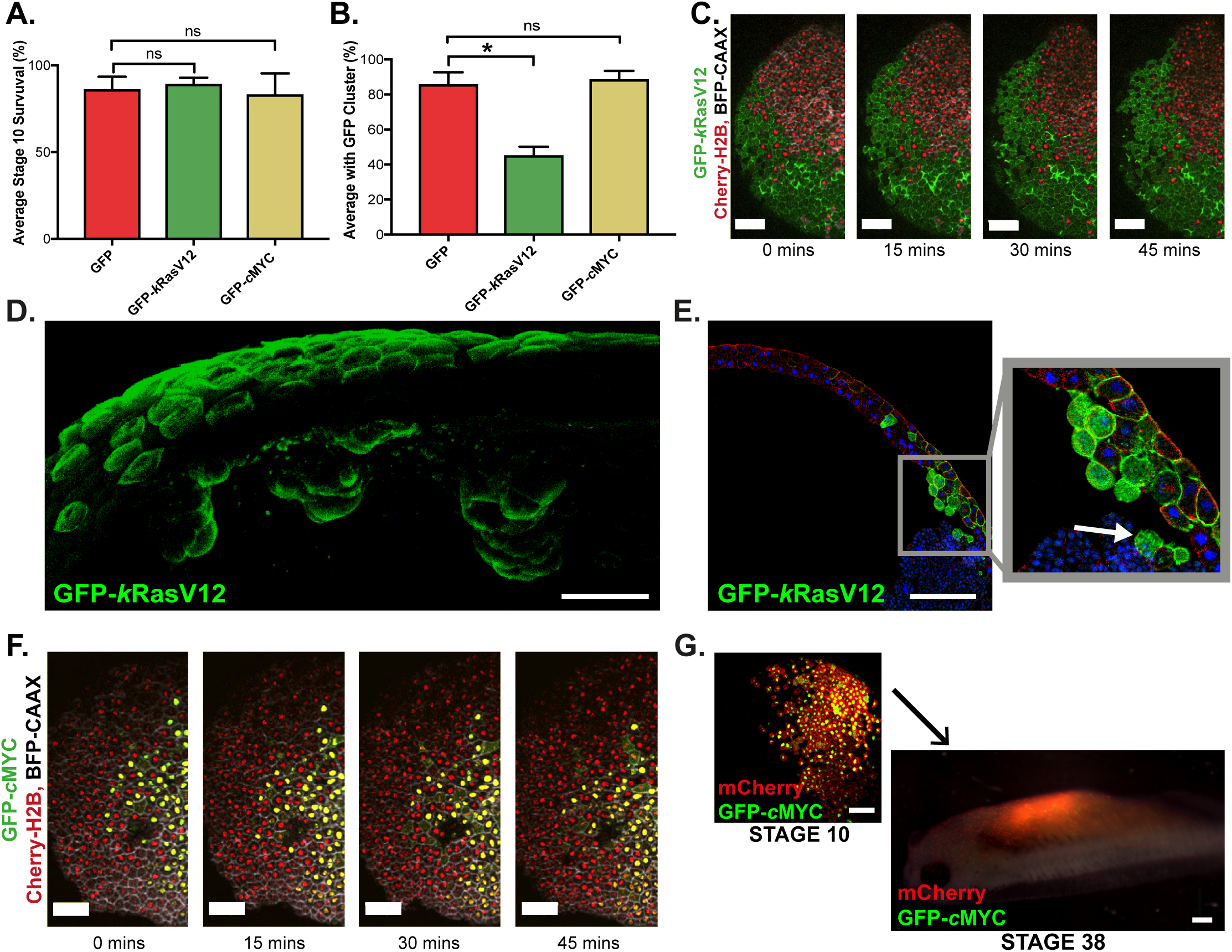
Further Characterisation of Oncogene-Expressing Cell Clusters in *Xenopus laevis*. **A.** Bar chart shows the average percentage of embryos, injected at the 32-cell stage, alive at stage 10. Kruskal-Wallis test: p>0.9999 for both, n=3 clutches of embryos. Error bars show SEM. **B.** Bar chart shows the average percentage of surviving stage 10 embryos, injected at the 32-cell stage, that have a GFP-positive cluster in the superficial animal cap layer. Kruskal-Wallis test: *p=0.0222, n=3 clutches of embryos. Error bars show SEM. **C.** Stills from a representative confocal microscopy time-lapse of a *Xenopus* embryo at early gastrula stage 10, with a GFP-*k*Ras^V12^ cell cluster in the superficial animal cap layer. No apical extrusion or apoptosis was observed in either the GFP-*k*Ras^V12^ clusters or the surrounding wild-type cells. **D.** Confocal microscopy image shows an embryo, where GFP-*k*Ras^V12^ mRNA was injected into a single cell at the 32-cell stage, that was fixed at stage 10, bisected and immunostained for GFP (green). **E.** Confocal microscopy image shows an embryo, where GFP-*k*Ras^V12^ mRNA was injected into a single cell at the 32-cell stage, that was fixed at stage 10, cryosectioned and immunostained for GFP (green), tubulin (red) and DAPI (blue). Arrows highlight cells that have lost cell-cell junctions and are no longer attached to the animal cap. **F.** Stills from a representative confocal microscopy time-lapse of a *Xenopus* embryo at early gastrula stage 10, with a GFP-*c*MYC cell cluster in the superficial animal cap layer. No apical extrusion or apoptosis was observed in either the GFP-*c*MYC clusters or the surrounding wild-type cells. **G.** Microscopy images show a representative embryo at stage 10 and stage 38 that was co-injected with GFP-*c*MYC and mCherry mRNA at the 32-cell stage. Anterior is towards the right. Scale bars represent 100µm in all panels, except D and G stage 38 where they represent 50µm and 500µm, respectively.

**Figure S2:**
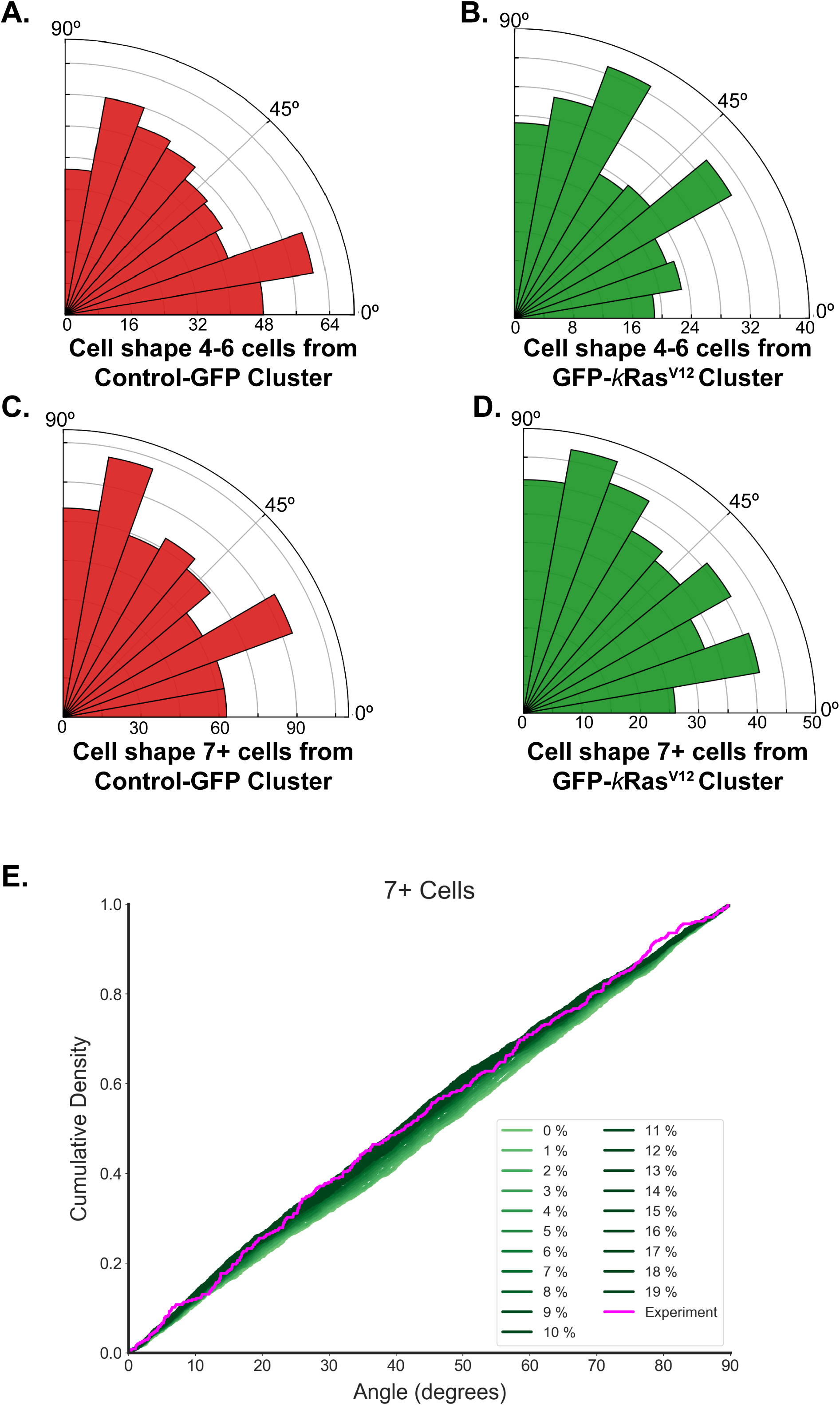
The Change in Wild-Type Cell Shape is a Localised Affect. **A-D.** Rose histograms show the orientation of wild-type cells long-axes 4-6 cells (A and B) and 7 or more (C and D) cells from GFP-control (A and C) or GFP-*k*Ras^V12^ (B and D) cell clusters, relative to the cluster, with the total number of cell divisions that were analysed across all embryos each data group in 10° bins. Kruskal-Wallis test: 4-6 cells: p=0.1572, n=433 cells from 5 GFP-control embryos and 240 cells from 5 GFP-*k*Ras^V12^ embryos. 7+ cells: p>0.9999, n=690 cells from 5 GFP-control embryos and 344 cells from 4 GFP-*k*Ras^V12^ embryos. E Cumulative distributions of cell shape orientation relative to cluster, 7+ cells from the cluster edge, comparing experiments (magenta) and simulations (green). Ras clusters were simulated with varying degrees of increased cortical contractility, Γ.

**Figure S3:**
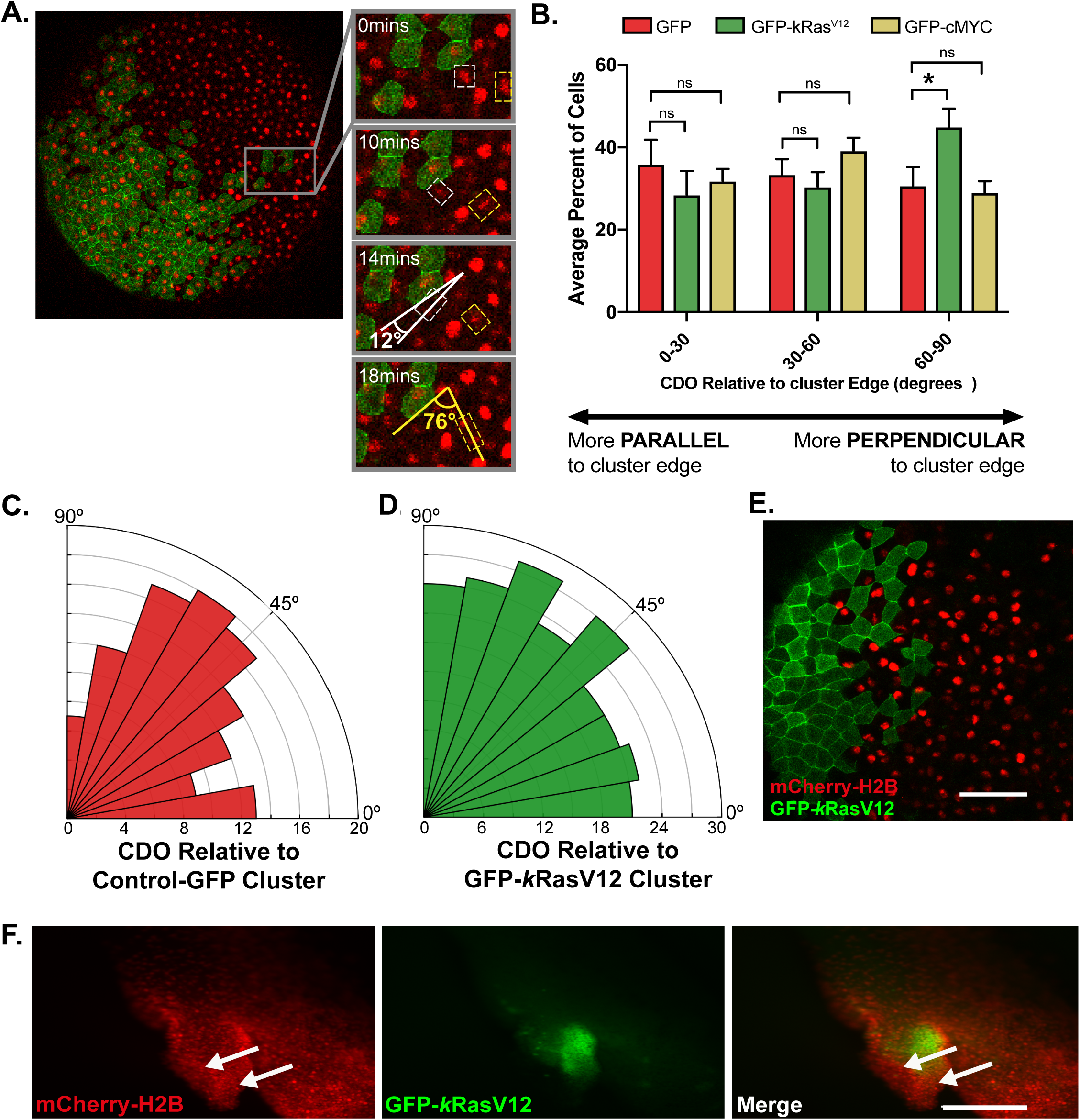
Further Characterisation of Wild-Type Cell Behaviour in *Xenopus laevis* Embryos with Oncogene-Expressing Cell Clusters. **A** Stills from a confocal microcopy time-lapse show the quantification of cell division orientation within the epithelial plane, relative to a GFP-expressing cluster. An angle of <u>90° indicates a division perpendicular</u> to the border of the cluster, whereas an angle of <u>0° indicates a division parallel</u> to it. **B** Bar chart shows the average percentage of cell divisions in 30° bins that occurred in wild-type cells up to 3 cells from GFP, GFP-*k*Ras^V12^ or GFP-*c*MYC clusters. Kruskal-Wallis test *p=0.0263; n=8 GFP, 9 GFP-*k*Ras^V12^ and 9 GFP-*c*MYC embryos. Error bars show SEM. **C-D.** Rose histograms show cell division orientation, relative to the cluster edge, of wild-type cells 7+ cells from (C) control-GFP or (D) GFP-*k*Ras^V12^ clusters, with the total number of cell divisions analysed across all embryos in each data group in 10° bins. Kruskal-Wallis test: p>0.9999, n=120 divisions from 8 GFP-control embryos and 212 divisions from 11 GFP-*k*Ras^V12^ embryos. **E.** Confocal microscopy image of a stage 10 *Xenopus* embryo injected with GFP-*k*Ras^V12^ (green) mRNA in a single cell at the 32-cell stage; mCherry-H2B (red) mRNA was then injected into neighbouring cells at the 32-cell stage. Scale bar is 100 μm. **F.** Microscopy image shows a representative ITLS, in a stage 38 embryo, that had a GFP-*k*Ras^V12^ cluster and wild-type cells labelled with cherry-H2B at stage 10. Arrows highlight wild-type tissue contributing to ITLS. Anterior is towards the right. Scale bar is 500 μm.

**Figure S4:**
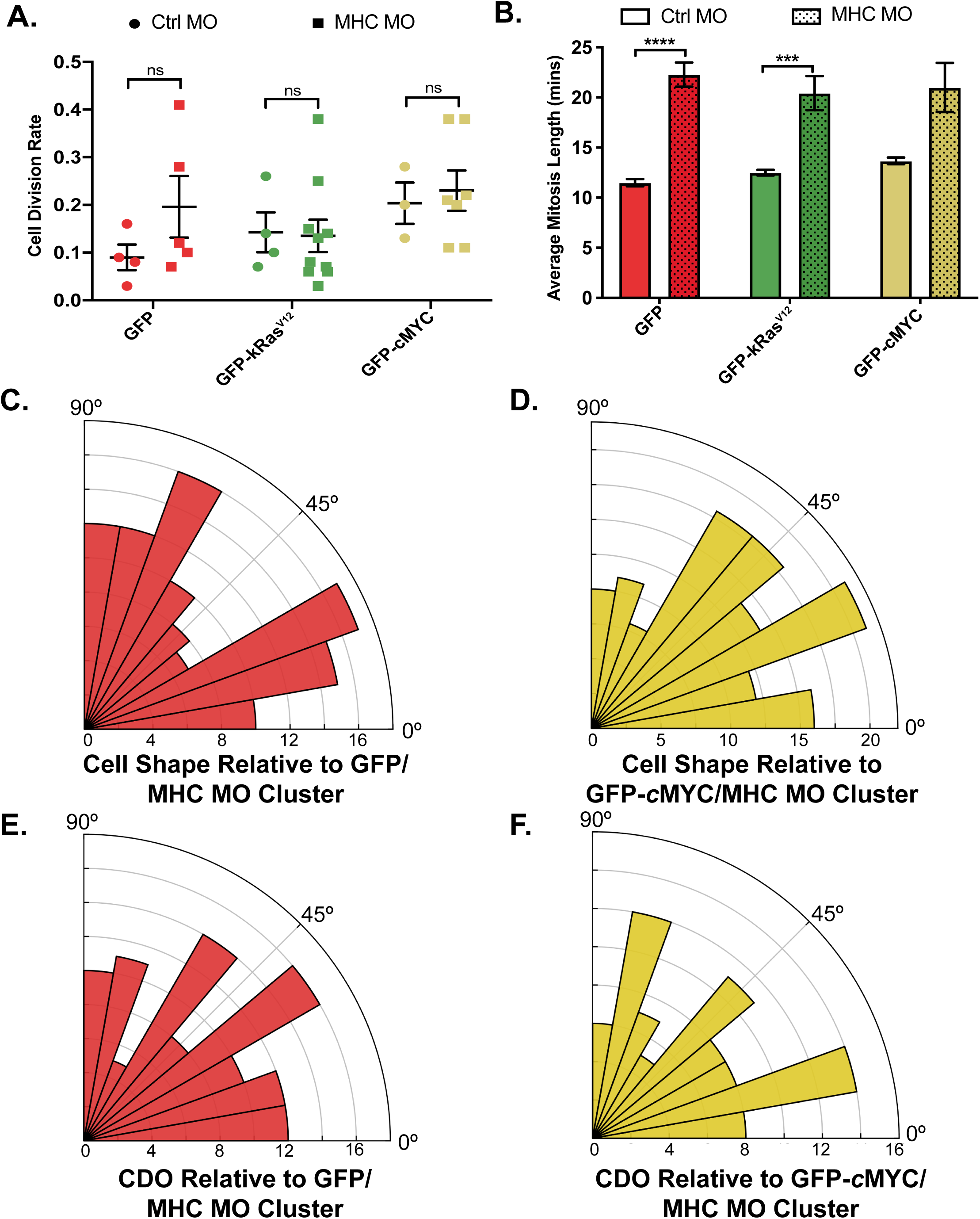
Further Characterisation of the Depletion of Myosin ll in Oncogene-Expressing Cell Clusters. **A.** Dot plot shows the average percentage of cells per minute that divided in clusters that were co-injected with GFP, GFP-*k*Ras^V12^ or GFP-*c*MYC mRNA and either control morpholino (Ctrl MO) or myosin heavy chain llB morpholino (MHC MO). GFP: Kruskal-Wallis test: n=4 GFP/Ctrl MO embryos, 5 GFP/MHC MO, 5 *k*Ras^V12^/Ctrl MO and 7 *k*Ras^V12^/MHC MO, 3 GFP-*c*MYC/Ctrl MO and 4 GFP-*c*MYC/Ctrl MO embryos. Error bars are SEM. **B.** Bar chart shows the average number of minutes between nuclear envelope breakdown and the separation of daughter nuclei in anaphase. Kruskal-Wallis test: ****p<0.0001, ***p=0.0007 n=12 GFP/Ctrl MO cells, 7 GFP/MHC MO, 12 *k*Ras^V12^/Ctrl MO, 9 *k*Ras^V12^/MHC MO, 12 GFP-*c*MYC/Ctrl MO and 4 GFP-*c*MYC/Ctrl MO. Error bars are SEM. **C-D.** Rose histograms show the orientation of wild-type cell long-axes up to 3 cells or from myosin ll deficient (C) GFP clusters or (D) GFP-*c*MYC clusters, relative to the cluster, with the total number of cells that were analysed across all embryos each data group in 10° bins. Kruskal-Wallis test performed against GFP/Ctrl MO shown in Figure 4G: GFP/Ctrl MO vs GFP/MHC MO p>0.9999, GFP/Ctrl MO vs GFP-*c*MYC/MHC MO p=0.0803, n=325 cells from 6 GFP/Ctrl MO embryos, 107 cells from 7 GFP/MHC MO embryos and 128 cells from 8 GFP-*c*MYC/MHC MO embryos. **E-F.** Rose histograms show cell division orientation of wild-type cells up to 6 cells from myosin ll deficient (E) GFP clusters or (F) GFP-*c*MYC clusters, relative to the cluster, with the total number of cell divisions that were analysed across all embryos each data group in 10° bins. Kruskal-Wallis test performed against GFP/Ctrl MO shown in Figure 4H: GFP/Ctrl MO vs GFP/MHC MO p>0.9999, GFP/Ctrl MO vs GFP-*c*MYC/MHC MO p>0.9999, n=58 divisions from 6 GFP/Ctrl MO embryos, n=99 divisions from 7 GFP/MHC MO embryos and 80 divisions from 8 GFP-*c*MYC MHC MO embryos.

## Notes

### Competing Interest Statement

The authors have declared no competing interest.

